# RNA polymerases in strict endosymbiont bacteria with extreme genome reduction show distinct erosions that could result in limited and differential promoters’ recognition

**DOI:** 10.1101/2020.09.07.285775

**Authors:** Cynthia Paola Rangel-Chávez, Edgardo Galán-Vásquez, Azucena Pescador-Tapia, Luis Delaye, Agustino Martínez-Antonio

## Abstract

Strict endosymbiont bacteria with high degree of genome reduction retain smaller proteins and, in certain cases, lack complete functional domains compared to their free-living counterparts. Until now, the mechanisms underlying these genetic reductions are not well understood. However, it is thought that, in order to compensate for gene reduction, somehow hosts take over those vital functions that endosymbionts cannot perform. In the present study, the conservation of RNA polymerases, the essential machinery for gene expression, is analysed in bacteria with extreme genome reductions. For this purpose, comparative genomics, phylogenetic analysis and three-dimensional models of RNA polymerase subunits were done over four lineages of endosymbiotic proteobacteria with the smallest genomes known to date. Amino acids under positive selection in the α subunit and loss of motifs in other subunits of RNA polymerase were observed. According to three-dimensional models, sites under positive selection might compensate the loss of motifs in α subunit. In addition, variations in the σ subunit were identified, some of them already studied in *E. coli* as a result of random mutagenesis. Amino acid changes in RNA polymerase suggest a possible modification in the binding specificity of the canonical −10 box (TATAAT) in some of these organisms. Furthermore, the β-flap helix domain is absent in some *Hodgkinia* strains, as observed in RNA pol II of Archaea, thus lacking the capacity to bind to the −35 box. Here, we propose several RNA polymerases models for endosymbiont bacteria with extremely reduced genomes. Evidence suggests that RNA polymerases of each endosymbiont bacteria follow a unique evolutionary path, without necessarily following the same path as a lineage, this is probably influenced by the intimate interactions sustained with other endosymbionts and its hosts.

## Introduction

Until 2006, there was a scientific consensus indicating that the minimum quantity of genes to support life would be around 500, even for tiny organisms like bacteria. This view changed soon after that date, when genomes of strict endosymbiotic bacteria began to be reported, and it was found that several genomes contained less than 500 genes, with some extreme cases reporting on genomes as small as 200 genes [1]. Transcriptome analysis of *Buchnera,* an obligate endosymbiont that has lost the ability to persist without its host, provided evidence indicating that, unlike *E. coli,* its genome has a limited ability to respond to environment fluctuations [2], which possibly indicates that gene expression in obligate endosymbiont bacteria is somehow stable and active at basal levels. According to this, the density of promoter-like signals, characteristic of free-living bacteria is not present in organisms exhibiting extreme genome-reduction [3].

DNA transcription is an essential molecular process through which organisms decode genetic information to cellular functions [4]. RNA polymerase (RNAP) is an enzyme responsible for the transcription of DNA to RNA, which consists of a multi-subunit protein complex present in all living organisms from bacteria to eukaryotes [5]. In bacteria, RNAP is responsible for the synthesis of all types of RNAs, including messenger RNAs, ribosomal RNAs, transference RNAs, and small RNAs. In free-living bacteria, RNAP consists of six subunits (α_2_ββ’ωσ), encoded by five different genes (the two α_2_ subunits are encoded by the same gene), the specific assembly of these proteins constitutes a holoenzyme with a molecular mass around 400 kDa. Previous studies on *Escherichia coli* found that the ω subunit is not essential for RNAP activity [6, 7], it was later found that ω subunit is absent in all the endosymbionts here analysed.

The rest of the subunits considered as essential core components of RNAP (α_2_, β and β’ subunits) are well conserved in bacteria [5, 8]. This RNAP core is catalytically active (transcribing DNA to RNA) but unable to initiate DNA transcription by itself. For transcription to be initiated, the core enzyme must bind to an additional subunit called sigma factor (σ). The σ is responsible for recognizing and binding to specific sites of transcription initiation, formally known as promoters; binding of sigma factors enables RNAP to initiate transcription [8]. Once transcription is initiated, and after the synthesis of a short fragment of RNA, σ is released and the core protein complex continues transcribing until it reaches a transcription terminator. Since σ is responsible of binding and discriminating among gene’s promoters, it is common to find different types (variants) of σ, in such a way that we can classify them in two evolutionary families: one of them called the σ^54^ family, that normally has one single gene per genome; and the σ^70^ family, that usually has several genes/members per genome (from 1 to 60 in the largest bacterial genomes). One member of the σ^70^ family is also known as the “housekeeping σ factor”, which is an essential gene present in all bacteria [9]. There are seven sigma factors encoded in the *E. coli* genome, six of them correspond to the σ^70^ family (σ^70^, σ^38^, σ^32^, σ^28^, σ^24^ and σ^19^), and the remaining one corresponds to σ^54^. In a previous work we analysed around 8500 promoters annotated in the *E. coli* genome and proposed the architecture of transcription units and the consensus sequences of the promoters for each sigma factor [10].

Endosymbionts live inside other organisms in a relationship estimated to be several million years old [11]. Strict endosymbiotic bacteria have lost most of their genes and large fragments of proteins due to a global process known as genome reduction [10, 6]. This dramatic process can be illustrated with the comparison of the *E. coli* genome, with approximately 4.5 thousand genes, and their relative endosymbiotic bacterium *Candidatus Carsonella ruddii,* which has only retained around 150 genes [12, 13]. Almost all genes found in *C. ruddii* are considerably shorter than their free-living orthologues. Furthermore, genes encoding proteins with multiple domains in strict endosymbionts commonly have lost some of these domains, which in some cases have been identified as essential for its activity in free-living bacteria [14, 15]. Endosymbiotic bacteria with extreme genome reduction a single copy of σ^70^ is maintained and none σ^54^ and in a previous work, we identified that obligate endosymbionts have lost all the transcription factors that interact with the promoters and RNAPs to activate or inhibit the transcription process [16]. Additionally, partial loss of the subunits α and σ have been reported previously in *Hodgkinia* sp. and *Carsonella ruddii*, respectively [15, 17], both subunits are essential for the proper operation of RNAP in free-living bacteria.

The phenomenon of gene erosion, including the transcriptional machinery in endosymbionts with extremely reduced genomes, raises questions concerning how gene transcription takes place in these bacteria. For instance: if the loss of RNAP subunits is happening in the same way in these bacteria, how is that some domains are dispensable in obligate endosymbionts while being essential in free-living bacteria? And, if we have elements to speculate, how the transcription process has been adapted in endosymbiont bacteria with extremely reduced. Here, we address some of these questions from the perspective of comparative genomics. We investigated the evolution of RNAP subunits in four bacterial groups of endosymbionts exhibiting extreme genome-reduction, all of them belonging to the proteobacteria phylum.

## Materials and Methods

### Dataset selection

The genetic information for 37 genomes of endosymbionts belonging to four lineages of proteobacteria with characteristics of extreme genome reduction were obtained from the NCBI database (http://www.ncbi.nlm.nih.gov/). These are *Candidatus Hodgkinia cicadicola,* that belongs to α-proteobacteria; *Candidatus Tremblaya phenacola*, *Candidatus Tremblaya princeps,* and *Candidatus Nasuia deltocephalinicola,* which belong to β-proteobacteria; and *Candidatus Carsonella ruddii,* that belongs to Gamma-proteobacteria. The genomes of these bacteria have been sequenced completely. Further characteristics of these genomes are provided in the supplementary material S1 Table. From these, in order to study the most reduced RNAP proteins (see below), we selected 16 bacterial endosymbionts genomes displayed the largest modifications at the protein domain level (Table 1).

**Table 1.** Genomic characteristics of the analysed endosymbionts compared to *E. coli*.

| Organism name | Lifestyle | Genome size* | Genes* | %GC | Reference |
| --- | --- | --- | --- | --- | --- |
| *Escherichia coli* k-12 MG1655 | Free-living | 4.63897 | 4405 | 50.8 | [13] |
| $\alpha$ proteobacteria | | | | | |
| *Candidatus Hodgkinia cicadicola* TETUND1 | Endosymbiont | 0.133698 | 121 | 46.8 | [18] |
| *Candidatus Hodgkinia cicadicola* TETUND2 | Endosymbiont | 0.14057 | 140 | 46.2 | [18] |
| *Candidatus Hodgkinia cicadicola* Tetuln | Endosymbiont | 0.14057 | 140 | 45.4 | [18] |
| *Candidatus Hodgkinia cicadicola* Dsem | Endosymbiont | 0.143795 | 169 | 58.4 | [19] |
| $\beta$ proteobacteria | | | | | |
| *Candidatus Tremblaya phenacola* PAVE | Endosymbiont | 0.1715 | 183 | 42.1 | [20] |
| *Candidatus Tremblaya princeps* PCIT | Endosymbiont | 0.138927 | 172 | 58.8 | [21] |
| *Candidatus Tremblaya princeps* PCVAL | Endosymbiont | 0.138931 | 174 | 58.8 | [22] |
| *Candidatus Nasuia deltocephalinicola* str. NAS-ALF | Endosymbiont | 0.112091 | 137 | 17.1 | [23] |
| *Candidatus Nasuia deltocephalinicola* str. PUNC | Endosymbiont | 0.112031 | 137 | 16.6 | [24] |
| Gamma proteobacteria |  |  |  |  |  |
| *Candidatus Carsonella ruddii* HT isolate Thao2000 | Endosymbiont | 0.157543 | 166 | 14.6 | [14] |
| *Candidatus Carsonella ruddii* PV | Endosymbiont | 0.159662 | 175 | 16.6 | [25] |
| *Candidatus Carsonella ruddii* PC isolate NHV | Endosymbiont | 0.159923 | 174 | 15.6 | [14] |
| *Candidatus Carsonella ruddii*<br>CS isolate Thao2000 | Endosymbiont | 0.162504 | 181 | 14.0 | [14] |
| *Candidatus Carsonella ruddii*<br>CE isolate Thao2000 | Endosymbiont | 0.162589 | 182 | 14.0 | [14] |
| *Candidatus Carsonella ruddii*<br>HC isolate Thao2000 | Endosymbiont | 0.166163 | 182 | 14.2 | [14] |
| *Candidatus Carsonella ruddii*<br>DC | Endosymbiont | 0.174014 | 196 | 17.6 | [26] |

\*Genome size in Mega base pairs, \*\*Open reading frame.

### Comparative analysis

The amino acid sequences for each RNAP subunit were aligned against their corresponding *E. coli* ortholog. For simplicity, the amino acid positions will hereinafter refer to its location in the corresponding *E. coli* protein or subunit, accompanied by the abbreviation “ECO”. For each endosymbiont, we used the protein sequences annotated for each RNAP subunit and made a comparative analysis to define domains and amino acid sites. The following tools were used for this purpose: Pfam database [31], the proteins superfamily classification database [32], the NCBI’s conserved domains search [33] and, additionally, we did a bibliographic research to gather relevant information regarding the sites and regions of each of the RNAP subunits and their functionality.

### Structural Analysis

To analyse protein structures, we first recreated 3D structural models for each RNAP subunit using the I-TASSER server [34]. For a given sequence, TASSER generates 3D models by collecting high-score structural templates from PDB, with full-length atomic models constructed by iterative template-based fragment assembly simulations. The resulting structures were compared with the crystal structure of the holoenzyme RNAP of ECO (4YLP) [35]. Graphic representations for each structure were prepared with the PyMOL Molecular Graphic System software version 1.3 [36].

To investigate possible dimer association of α subunit monomers of *Hodgkinia* TETUND2 (this strain has lost a fragment of the α subunit amino terminal and it is involved in dimer formation), we used the ClusPro v.2.0 tool [37–40]. This is an automatic protein docking tool based on CAPRI (Critical Assessment of Predicted Interactions) [38, 41]. Three models of α dimer subunits with different cluster sizes were obtained through ClusPro and were used to map sites under putative positive selection and for evaluating free-energy changes in protein-protein binding. With this strategy, we recreated single mutations along the amino acid residues positively selected using the BindProfX server [42]. We exchanged the putative positive selected amino acid on the dimers predicted by the amino acid from different strains of *Hodgkinia* and determined the changes in protein binding affinity. The binding affinities between each pair of proteins were measured as Gibbs free-energy change ΔG=G(complex)-G(monomers) when two monomers form a complex, the more negative a ΔG is, the more stable the complex is. The effect of mutations on binding affinity were measured by the differences on free-energy changes between the mutant and the wild type ΔΔG_wt->mut_=ΔGmut−ΔGwt. A strongly favourable mutation was considered to have ΔΔG≤−1kcal/mol.

### Selective pressure analysis

To study the putative mechanisms of molecular evolution of the RNAP subunits, first we need to know the selective pressure acting on each of the subunits encoding for these genes. With this purpose, we performed D_N_/D_S_ tests on gene/protein sequence, calculated as the ratio D_N_/D_S_ of the number of non-synonymous substitutions per non-synonymous sites (D_N_) divided by the number of synonymous substitutions per synonymous sites (D_S_). In such a way that D_N_/D_S_ could take one of three values, i) if non-synonymous mutations are deleterious, *i.e.* purifying selection will reduce their fixation rate and D_N_/D_S_ value will be less than 1; ii) if non-synonymous mutations are advantageous they will be fixed at a higher rate than synonymous mutations, and consequently the D_N_/D_S_ value will be greater than 1; and iii) if the D_N_/D_S_ ratio is equal to 1, it indicates that the evolution is neutral [43]. Selection analysis was performed for RNAP subunits in each group of bacteria.

Each set of genes was implemented in the codeml from PAML v.4.6 package [44], which uses an alignment by codons that was created using the Codon Alignment PAL2NAL v.2.1.0 program [45]. Phylogenetic trees were built using PhyML [46], the set of amino acids sequences of each subunit were first aligned with T-Coffee [47] and the informative blocks were recovered using Gblocks [48]. Two approaches, branch and branch-site were used to identify genes and amino acids under directional positive selection. For branches, three models were used; “M0” one-ratio model (D_N_/D_S0_), Free model (D_N_/D_S1_) and two-ratio model D_N_/D_S2_. The D_N_/D_S0_ model assumes the same D_N_/D_S_ ratio for all the branches, the D_N_/D_S1_ assumes an independent D_N_/D_S_ for each branch and the D_N_/D_S2_ assumes that the interest branch (foreground branch) has a D_N_/D_S2_ ratio different than the background ratio [49].

The level of significance for the Likelihood Ratio Test (LRT) was estimated using the *x*^^2^^ distribution with degrees of freedom (df), number equal to the difference of the number of parameters between the models and the calculated statistic was defined as twice the difference of log-likelihood between the models (2ΔlnL= 2[lnL_1_−lnL_0_]; where L_1_ and L_0_ are the likelihoods for the alternative and null models, respectively) [50]. One ratio and free ratio models were compared in order to know whether D_N_/D_S_ were different among the lineages, while one ratio and two-ratio models were compared in order to examine whether the lineage of interest has a different ratio compared to other lineages.

To detect positive or negative selection in specific lineages displaying D_N_/D_S_ values greater or smaller than one, we approached through other models, in which model two and the D_N_/D_S_ ratio were fixed to 1 (D_N_/D_S=1_), 0.2(D_N_/D_S=0.2_) and 1.2(D_N_/D_S=1.2_) for the foreground branch. First, we compared the models D_N_/D_S2_ against the D_N_/D_S=1_, where the null hypothesis is that in which the models are not significantly different. If the null hypothesis is rejected (p<0.05) and the two-ratio model is greater than 1, it indicates the possibility of positive selection in the foreground, otherwise, if the two-ratio model estimate is smaller than 1, it is indicative of negative selection. On the other hand, if the null hypothesis is accepted, it is evidence that the foreground branch is under neutral selection (without selection). Additionally, we compared the models D_N_/D_S2_ with the D_N_/D_S=0.2_, where the null hypothesis is that in which the models are not significantly different. If the null hypothesis is rejected (p<0.05) and the two-ratio model estimate is greater than D_N_/D_S_ =0.2, it indicates a weaker negative selection, while a value smaller than D_N_/D_S_ =0.2 indicates a stronger negative selection. Finally, when D_N_/D_S2_ is greater than 1 and it is significantly different to D_N_/D_S=1_, it indicates positive selection. In order to get more evidence about possible positive selection we compared D_N_/D_S2_ against D_N_/D_S=1.2_, the null hypothesis was defined as non-significant difference between D_N_/D_S2_ and D_N_/D_S=1.2_, acceptance of the null hypothesis indicates that the foreground branch is possibly under positive selection, while a rejection indicates that the foreground branch might be subjected to a relaxed selection.

To identify individual codons under positive selection along specific branches, we performed a branch-site test for positive selection [50]. In these models, positive selection was allowed on a specific, “foreground” branch, and the LRTs (df=1) were performed against null models that assume no positive selection is happening. This test has four classes of sites: 0, 1, 2a and 2b; for the site classes 0 and 1, all codons are under purifying selection (0< D_N_/D_S0_<1) and neutral evolution (D_N_/D_S1_=1) for all branches, respectively. For the sites in classes 2a and 2b, positive selection is allowed on the foreground branches (D_N_/D_S2_>1), but the rest, the “background branches”, are under purifying selection (0<D_N_/D_S0_<1) and neutral evolution (D_N_/D_S1_=1), respectively. For the null model, D_N_/D_S2_ is fixed to 1. In our analysis, all the RNAP subunits in each endosymbiont group were tested that way. Phylogenies were reconstructed by testing each branch as foreground. We compared the two models using LRT and the significance between the models was assessed by calculating twice the log-likelihood difference following a x^2^ distribution, with a df number equal to the difference of the number of parameters between the models. Positive selected amino acid sites are identified based on Empirical Bayes and posterior probabilities were employed in codeml [51]. We did not test the *Nasuia* RNAP subunits, because there were only three sequenced strains but codeml requires at least 4 in order to get reliable results.

## Results and Discussions

To evaluate the putative evolution of RNAP’s in endosymbiotic bacteria exhibiting extreme genome reduction, we identified orthologous genes of the *E. coli* RNAP genes present in the 37 available genomes of endosymbiotic bacteria (S1 Table). These include 9 strains of *C. ruddii*, 17 strains of *H. cicadicola*, one strain of *T. phenacola*, 7 strains of *T. princeps,* and 3 strains of *N. deltocephalinicola*. Of those, 16 genomes displayed the largest modifications at the protein domain level. For this reason, we decided to describe the RNAP subunits of these 16 organisms (Table 1, Fig 1).

**Fig 1.**
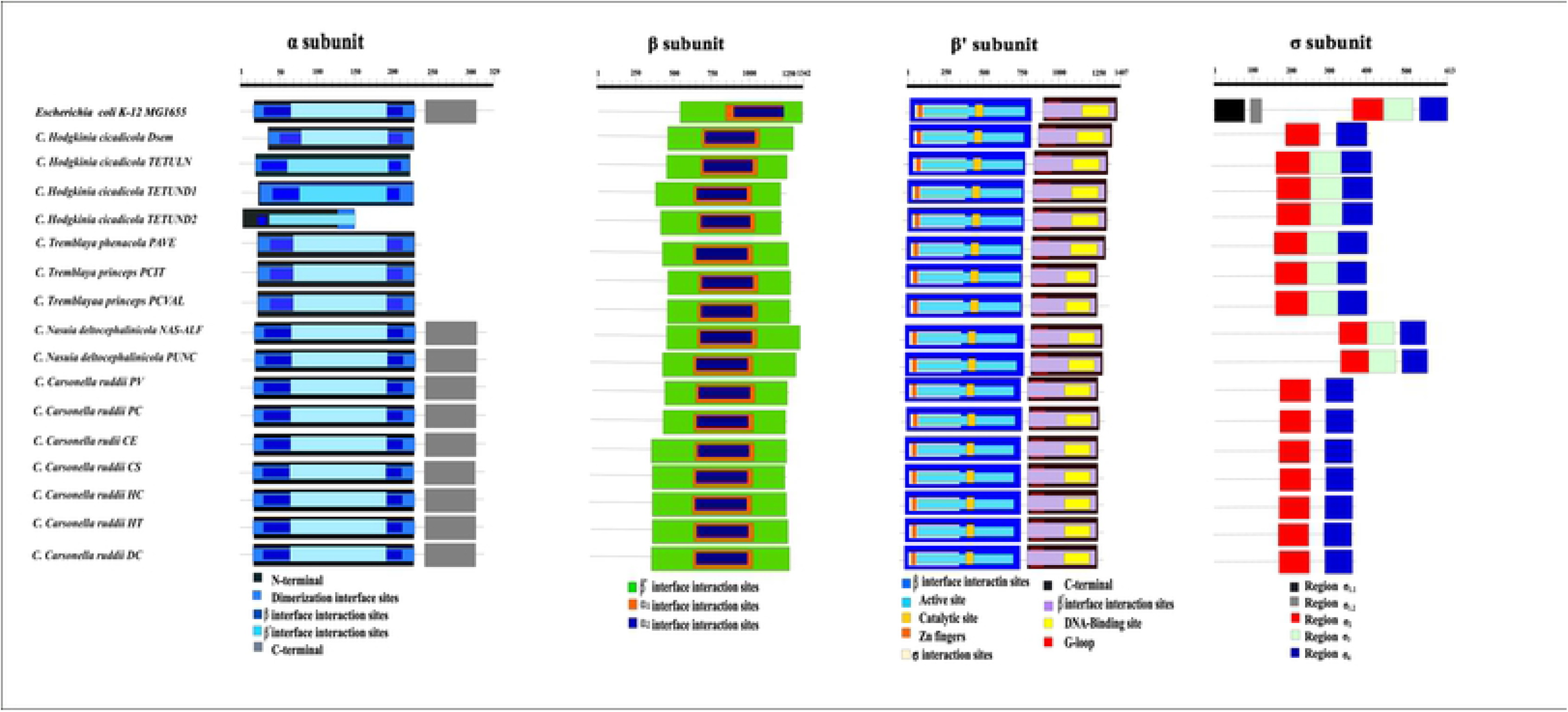
Conserved domains in RNAP subunits of endosymbionts compared to *E. coli*. Conservation of α subunit domains; black fragments represent N-terminal domains, light blue domains represent dimer interfaces, dark blue domains interact with the β subunit, turquoise domains interact with β’, and grey fragments represent the carboxy-terminal end. In the β subunit; green domains interact with β’, orange domains interact with α subunit 1, dark blue domains interact with α subunit 2. In the β’ subunit; light blue domains interact with β subunit, catalytic sites are shown in gold, sites interacting with sigma subunits in beige, zinc fingers in orange, active sites in turquoise, black fragments represent carboxy-terminal domains, β prime interface interaction sites are shown in purple, DNA binding sites in yellow, and G-loop domains in red. σ subunit domains; region 1 in black, region 1.2 in grey, region 2 in red, region 3 in mint, and region 4 in blue.

All bacteria conserve orthologous genes for each of the α, β, β’ subunits, as well as a gene coding for a σ subunit. On the other hand, the ω subunit, with in *E. coli* assists during the final step of RNAP core assembly by helping to bind the β’ subunit to the α2β sub-complex, it is no longer conserved in any of the analysed endosymbiotic bacteria. Previous studies revealed that ω is not an essential subunit in free-living *E. coli* and therefore it is not surprising that this gene has been lost in all these endosymbionts [15].

Previous works report that proteins in endosymbionts with reduced genomes tend to be shorter compared to free-living bacteria, which leads to the loss of protein regions [10, 14]. We calculate that the DNA sequences encoding for each of the RNAP subunits exhibit a reduction of 16% on average in comparison to those in *E. coli.* In addition, we identified that these genes have lost DNA regions that code for functional protein domains in *E. coli*. That is the case of the α subunit, which lacks the α-carboxy terminal domain in both Tremblaya and *Hodgkinia* strains as well as a part of the domain for dimer formation domain in *Hodgkinia* TETUND2 (Fig 1a). The β and β’ subunits conserve all the important functional domains (Fig 1b and Fig 1c) and sigma has lost regions 1 and 1.2 in all endosymbionts and region 3 in *Carsonella* strains (Fig 1d). In the next sections, we will describe in more detail the structure and variations found in each RNAP subunit of these endosymbionts.

### The α subunit

In *E. coli*, the gene *rpoA* codes for the RNAP α subunit, this protein has a molecular weight of ~37 kDa with two independently folded domains connected by two flexible linkers (α carboxy terminal and α amino terminal domains, corresponding to the grey and black boxes in Fig 1a, respectively) [48, 49, 50]. This subunit has three described biological functions: i) it initiates the assembly of the RNAP complex through the interaction of its amino-terminal domain (αNTD) with β and β’ subunits (Fig 2, black upper bar), ii) it participates in promoter recognition through the interaction of its carboxy-terminal domain (αCTD) (Fig 2, grey upper bar) with the DNA UP promoter element, and iii) its carboxy-terminal domain (αCTD) is also a target for the binding of a large number of transcription factors acting as transcriptional activators [52, 53, 54].

**Fig 2.**
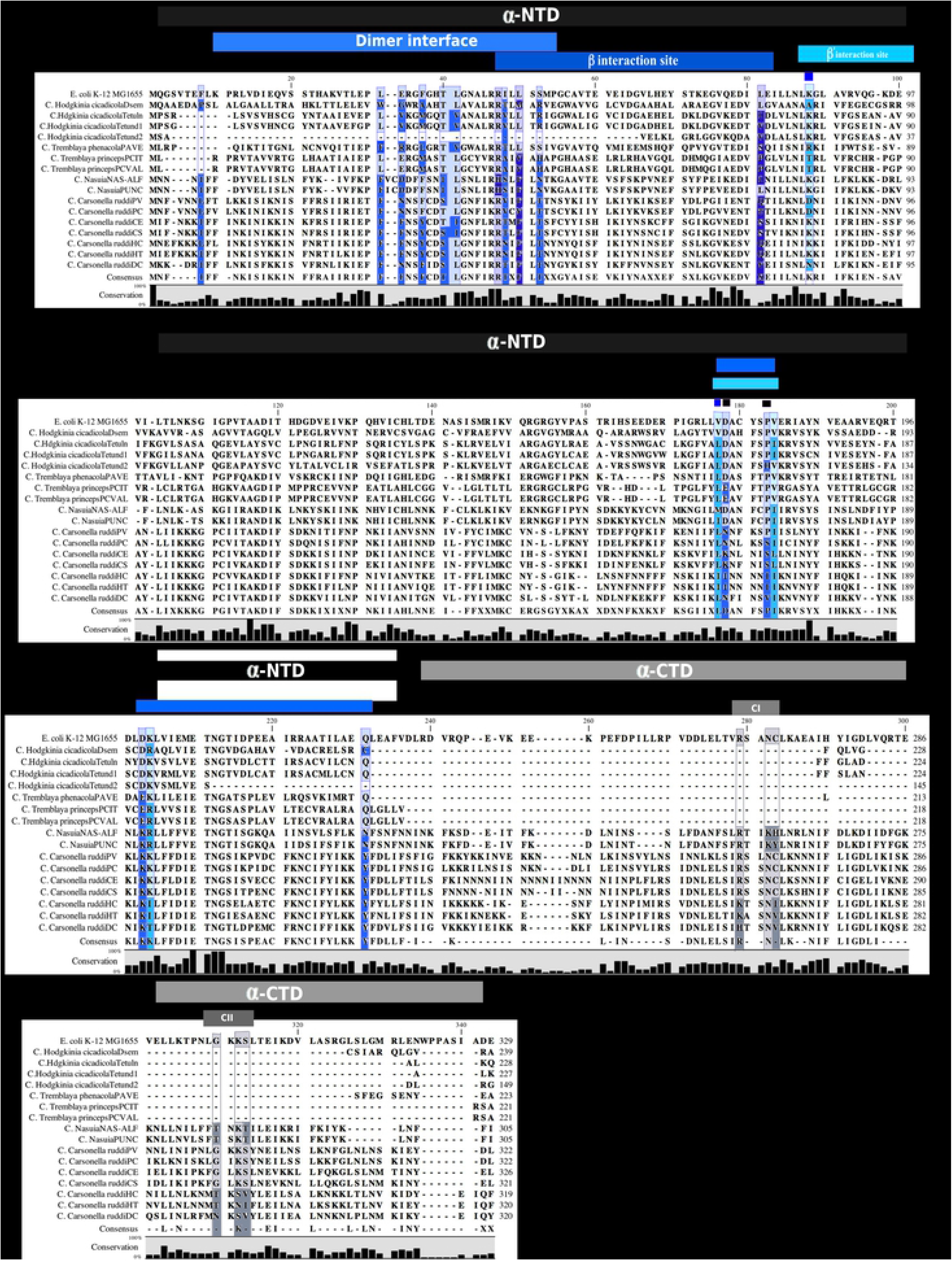
Functional domains and amino acid sites of the α-subunit. Black and grey upper bars on the amino acid’ alignment represent the domains corresponding to amino- and carboxy-terminal ends. Light blue represents the α subunits dimerization region, dark blue represents the region and the main site of interaction with β subunit, and turquoise represents the main sites of interaction with the β’ subunit. Those functional residues that differ from the reference organism (*E. coli)* are shown in darker backgrounds.

The αNTD domain in turn consists of two subdomains: subdomain 1 is called the dimerization domain, and subdomain 2 contains the interface sequences for the β and β’ subunits. These subdomains are well conserved among all the α subunits in prokaryotes, archaea and eukaryotes. The subdomain 1 consists of four anti-parallels β-sheets and two orthogonal α helices (H1 and H3). The subdomain 2 is conformed of one α helix and seven β strands in an antiparallel arrangement and are determinant for the interaction of α subunits with the β and β’ subunits [55, 56, 57].

The first step towards RNAP core formation is the homodimerization between two monomers of α subunits. This dimer is formed by the interaction of H1 and H3 helices; such interaction takes place in subdomain1 of each monomer. There are nine residues conserved in the αNTD, five of them are in the H1 and H3 helices (ECO: 35F, 38T, 39L, 46I, 227Q). Other three residues (ECO: 32E, 8F, 50S) are in the loop regions connecting the two α helices and the β sheets, and L31 is localized in the strand two of the C-terminal end (Fig 2, light blue sites).

Punctual mutations affecting residues 45R, 48L or 80E in *E. coli* have shown to prevent the binding of α_2_ to the ββ’ subunits (Fig 2, dark blue), but do not impede the homodimerization between α subunits. Furthermore, mutations affecting residues 86K, 173V, 180V or 200I (Fig 2, turquoise sites) in *E. coli* prevent the formation of α_2_ββ’ complex, but don’t do so for the α_2_ β complex; hence, these results suggest that these residues are important for the binding of α and β’ subunit [57].

In the case of *Carsonella* and *Nasuia* strains, their α subunits conserved all the functional domains as described in *E. coli*, while *Hodgkinia* and *Tremblaya* strains just conserved the αNTD (dimerization of β and β’ interaction domains) (Fig 2 light blue, dark blue and turquoise sites and S1 Fig). *In vitro* and *in vivo* experiments revealed that αNTD is essential for RNAP to achieve basal transcription [58, 59]. On the other hand, the αCTD is unnecessary for RNAP assembly and basal transcription but it is required for the interaction of DNA UP-promoter elements and transcription activators in *E. coli* [60, 61]. As mentioned, all these bacteria exhibiting extreme-genome reduction do not conserve any transcription factor in their genomes; and therefore, the absence of the αCTD domain would not impact the basal functionality of RNAP.

The loss of αCTD and other protein domains has also been reported in other studies conducted on bacteria, most of them belonging to the phylum *Parcubacteria,* which are ectosymbionts that live in mixed groups of bacterial communities, and therefore may be subject to a relaxed selection process [62]. The absence of αCTD is also common in some microalgae chloroplasts. In the case of genes encoded by chloroplasts, they are transcribed by two kinds of RNA polymerases, one of these is related to one-single protein phage RNA polymerase encoded in the nuclei genome of microalgae. The other RNAP is more related to the multi-subunit bacterial RNAP and is encoded by the in-chloroplast genome. The absence of αCTD in plant chloroplasts RNAP suggests that recent changes had happened in this clade [63].

A particular case is *H. cicadicola* TETUND 2, which partially conserves an αNTD, precisely located for its interaction with the β’ subunit and preserving a few amino acid residues for homodimerization (Fig 2, light blue sites). With the 3D predicted model of this subunit, we observed that the α subunit does not conserve the H1 and H3 helices to form the dimers interface (Fig 5a, white structures). The absence of these two α helices suggests a loss of capacity to form that dimer, which is essential for the RNAP core formation. *H. cicadicola* TETUND 2 and *H. cicadicola* TETUND 1 share the same host, but they are losing different genes. The *rpoA* gene is not the unique case of genes losing a large fragment of their protein products in *H. cicadicola* TETUND 2; the *dnaQ* gene that encodes the ε subunit of DNA polymerase III has also lost large fragments, in such a way that it is now considered as a non-functional gene in Tetund 2. A recent analysis reported additional cases of other *Hodgkinia* strains undergoing triple and sixfold coexistence with other *Hodgkinia* strains, which suggests gene complementation [17]. On the other hand, the genome of *Hodgkinia* doesn’t code for transporters or other proteins involved in cell envelope biosynthesis that could be mediating the exchange of proteins with other strains [18], hence, it is unclear how these bacteria maintain a functional metabolism.

### β and β’ subunits

The *rpoB* and *rpoC* genes encode the β and β’ subunits of RNAP, respectively. The β’ subunit forms a pincer, called “clamp”, and the β subunit constitutes the other pincer, giving place to a 27Å-wide internal channel, where the catalytic site of the RNAP enzyme is located. For the folding of RNAP, it is necessary for the β subunit to bind to the α and β’ subunits. Crystallographic patterns obtained from RNAP of different organisms suggests the participation of 24 (ECO: 810-1224) and 73 (ECO: 546-1340) amino acid residues in the β subunit is necessary for its binding to α and β’ subunits, respectively [64] (Fig 1b Green, orange and dark blue domains, respectively).

On the other hand, on β’ subunit, the interaction sites with the β subunit (at positions ECO: 917-1361, Fig 1c, light blue domain) and ω subunit (ECO: 907-1316) are located on the amino-terminal domain. Additionally, in the amino-terminal domain of the β’ subunit there are several functional residues that are typical of Zinc fingers (ECO: C70, C72, C85 and C88) (Fig 1c, orange region), involved in forming the catalytic site and giving stability to the RNAP during its interaction with DNA. The active site is formed by around 500 amino acids retaining two Mg^2+^ ions. However, only thirteen amino acid residues have been identified to be the most conserved ones (ECO: P121, R311, K334, R339, R346, R351, R425, A426, P427, T790, A791, G794 and Y795, Fig 1c, turquoise sites); they are adjacent to the catalytic site, which is composed by three aspartates (ECO: D460, D462 y D464, Fig 1c, golden sites) [65]. In the β’ C-terminal domain three polar residues (ECO: R1148, K1311 and E1327, Fig 1c, yellow region) and the G-loop domain (ECO: 1238-1250, Fig 1c, red region) form the cavity where the DNA fits and contact the RNAP.

β and β’ amino acid sequences are some of the most conserved among RNAP subunits in endosymbionts. Their critical role in RNAP complex formation may be the reason why these subunits are well conserved even in bacteria containing extremely reduced genomes (S2 Fig and S3 Fig).

Once the RNAP core is formed, σ and β’ bind together by means of a protein region contained in the β’ subunit, known as the β’ coiled-coil domain (ECO: 262-309, Fig 1c, beige region), which interacts with the σ_2_ domain of the σ factor (see below). Such binding puts the σ factor in contact with the −10 box, forming an open promoter complex. Different *in vitro* studies involving punctual mutations in the β’ coiled-coiled domain residues (ECO: R275Q, E295K and A302D) (Fig 3b, beige sites), resulting in a deficient holoenzyme formation and a subsequent lack of promoter specificity [66]. In addition, the β’-zipper region also interacts with the −10 box element of DNA promoters; this phenomenon largely involves the amino acids residues ECO: Y34 and R35, which form polar interactions with the polyphosphate of the nucleotide in position −17 of the promoter [45, 46].

**Fig 3.**
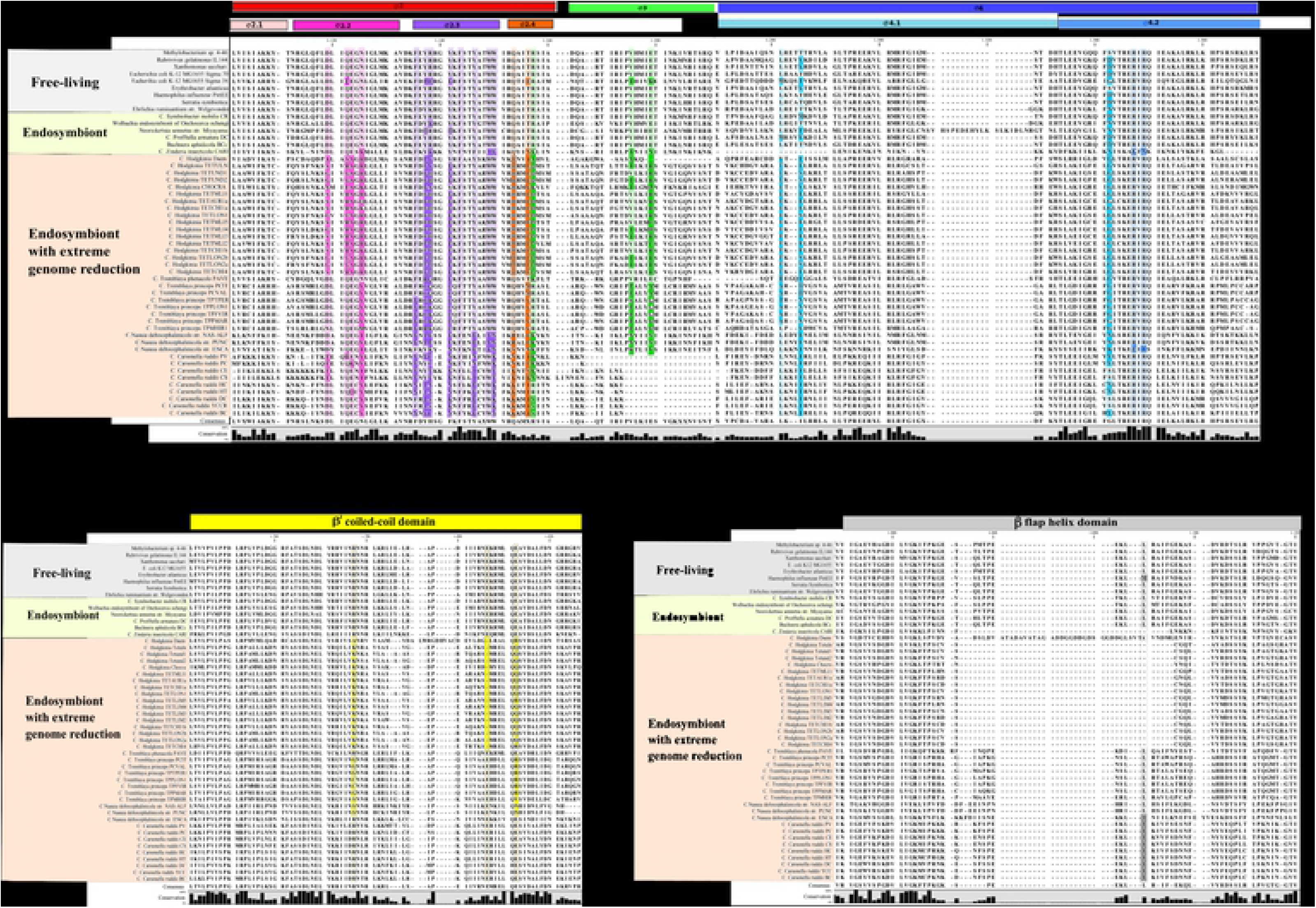
Sigma regions and sites involved in promoter recognition and binding. a) Sigma factor regions: σ_2_ (red), σ_2.1_ (pink), σ_2.2_ (magenta), σ_2.3_ (violet), σ_2.4_ (orange), σ_3_ (green) and, σ_4_ (dark blue), σ_4.1_ (cyan) and σ_4.2_ (light blue) b) Regions and important sites of the β’ subunit forming the β’-coiled coil domain (yellow) by interacting with a sigma factor c) Region and important sites in the β subunit to form the β-flap tip helix domain (grey).

Likewise, the β subunit binds to the σ factor through the β-flap helix domain (ECO: 889-898) (S2 Fig, grey sites) and the σ_4_ region of the σ factor (Fig 4c). The capacity of RNAP to adapt to variation in nucleotide distance between the promoter boxes (−10 and −35) depends on the aforementioned binding [67].

**Fig 4.**
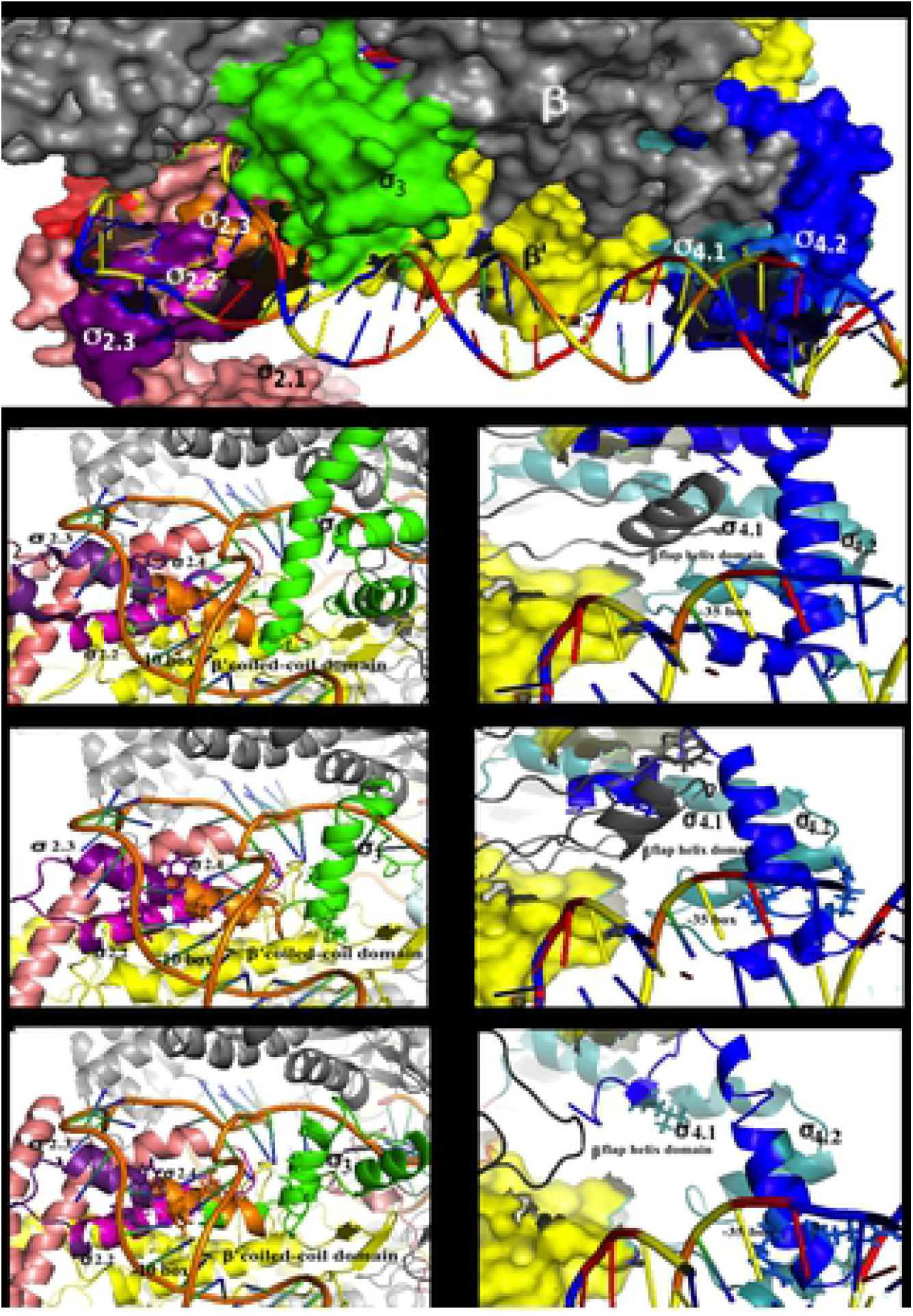
Structural comparison between *E. coli* and *Hodgkinia* sigma factors. a) Crystallography structure of *E. coli* RNA polymerase bound to a promoter (4YLP). The figures b, d and f show the structure of regions involved in the recognition and binding of the −10 box of *E. coli* (4YLP), *C. Hodgkinia* Dsem and TETUND2 promoters respectively. Figures c, e and g show the structure of the regions involved in the recognition and binding of the −35 box in *E. coli*, *C. Hodgkinia* TETUND2 and Dsem. The colours in the structures correspond to the colours in the amino acid alignment of Fig 3.

Except for *T. phenacola* PAVE and *C. ruddii* CE, which conserve these domains, the rest of endosymbionts display substitutions in at least one of the three important β’coiled-coil amino acids (Fig 3b yellow sites and S3 Fig). The β-flap helix domain is conserved in all endosymbionts here studied, including *Hodgkinia* Dsem. However, other *Hodgkinia’*s strains lack this domain (Fig 3c grey sites and S2 Fig). Mutants of the *E. coli* β subunit involving β-flap helix removal result in an inability of the σ subunit to bind the −35 box, without affecting the ability of the RNAP core to bind DNA. Furthermore, these mutants adequately recognize the −10 box and −10 extend promoter elements. The absence of β-flap domain has not been observed in bacteria previously; however, this condition is common in archaeal RNAPs. It has been proposed that the β subunit of archaeal RNAPs recognizes both the TATAAT Box (TB) and an accessory DNA motif located upstream from it [68]. This could suggest that with to exception of *Hodgkinia* Dsem, the RNAP of *Hodgkinia*’s strains cannot bind the −35 box, and instead it would only recognize the −10 box and the −10 extended promoter region during transcription initiation.

### σ *subunit*

Sigma factors (σ) are not permanent components of RNAPs core and are the last subunit to be associated with the RNAP complex to initiate DNA transcription. As mentioned, σ is needed for promoter recognition in *E. coli*. This subunit dissociates from the core enzyme once RNAP begins with the RNA synthesis process. In the case of *E. coli,* it encodes a total of seven sigma factors; however, the most essential one is the σ^70^ factor, encoded by the *rpoD* gene. σ^70^ is formed by four helical domains (σ_1_, σ_2_, σ_3_, and σ_4_; Fig 1c black, red, mint and blue boxes, respectively), each of them interacts with different promoter elements as well as with domains of the subunits forming the core enzyme. The σ_1.1_ subdomain, once inserted into the apo-holoenzyme, is located into the downstream channel, in contact with both sides of the RNA cleft [69]. σ_1.1_ prevents σ^70^ from interacting with the DNA strand in the absence of the RNAP core [70].

In the meanwhile, the σ_2_ domain (Fig 3a red sites) is the most conserved of the σ^70^ family. This region is divided into four sub regions: σ_2.1_ (Fig 3a, light pink bar), σ_2.2_ (Fig 3a, magenta bar), σ_2.3_ (Fig 3a, violet bar) and σ_2.4_ (Fig 3a, orange bar). While the σ_2.2_ contains the sites involved in the RNAP core binding (ECO: D403, Q406, E407 y N409) (Fig 3a, magenta sites), the σ_2.3 y_ σ_2.4_ participate in DNA melting and the recognition of the −10 box, respectively (Fig 3c). Likewise, the σ_2.3_ is characterized by the presence of seven conserved aromatic amino acids (ECO: F419, Y421, Y425, F427, Y430, W433 and W434) (Fig 3a, violet sites), and the replacement of ECO: Y425, Y430, W433 and W434 by alanine results in defects in DNA melting activity. In addition, previous studies showed that residues F419 and F427 play an important role for the precise folding of the α helices of the σ_2_ region. Furthermore, the specific recognition of −10 boxes has been attributed to the σ_2.4_ region (Fig 3a, orange bar). Multiple studies on σ^70^ have shown that the residues ECO: Q437 and T440 have implications in the recognition of the thymine in position −12 of the core promoter [71, 72]. On the other hand, the position −13 in the σ^70^ promoters is not conserved, and the substitution of residue ECO: 441R in σ has shown to be implicated in the recognition of nucleotides at this position. Amino acids involved in the DNA melting and recognition of the −10 box are exposed on the same face of the DNA helix. Deletion analyses have determined that the residues ECO: 361-390, comprising the most of σ_2_ region, seem to be necessary for RNAP correct functioning [72, 73].

The σ_3_ region (Fig 3a, green bar) interacts with the −10 extended promoter element. Additionally, the σ_3_ loop might have a role in stabilizing the short nascent DNA-RNA hybrids during the early stage of transcription initiation [74]. More specifically, this is the result of an interaction between the 5’-triphosphate of the nascent RNA with the σ_3.2_ loop of the σ factor [75].

Finally, the σ_4_ region (Fig 3a dark blue upper bar) is formed by two sub regions, σ_4.1_ and σ_4.2_ (Fig 3a, cyan and sky blue respectively), comprising four helices. The σ_4.1_ region interacts with the β-flap domain (Fig 4c) and, therefore, the distance between the σ_2.4_ and σ_4.2_ increases and allows their binding to the − 35 box of a promoter. Additionally, this region is also a point of contact with activators that bind upstream of the −35 box. The strength of the interaction between the β-flap domain and the σ_4.1_ region has effect in promoter specificity. As an example, one study reports that the binding of σ^38^ with the β-flap domain is stronger than with σ^70^, so the interaction of σ^38^ with the −35 box is stronger than that of σ^70^, with independence of the promoter sequence. Finally, in the σ_4.2_ region the third and fourth helices form a helix-turn-helix structure that interacts with the promoter at the −35 box, specifically, the amino acids R518 and R516 (Fig 3a, sky blue sites) recognize the guanine and cytosine in the −34 and −32 positions, respectively [73].

The σ subunit is the one that exhibits the most differentiated preservation pattern among endosymbionts with extreme genome reduction. In the case of *T. phenacola* PAVE, it conserves the σ_2_ and σ_4_ regions. On the other site, *Hodgkinia, N. deltocephalinicola* and *C. ruddii* only conserve the σ_4_ region and to a lesser extent the σ_2_ (Fig 3a). The main variations were observed in the σ_2_ region, where the residues that participate in the interaction with the −10 box and define the promoter specificity are located, particularly in the σ_2.2_ and σ_2.4_ sub regions.

For the σ_2.2_ sub-region, *Hodgkinias* exhibit substitutions at all four residues involved in the interaction with the β’ coiled-coil domain. The 17 strains have the same substitutions for the sites ECO E407A and 14 of them ECO N409R, while each one retains distinct amino acids at the positions ECO: 403 and ECO 406 (Fig 3a, magenta sites). These variations in the σ_2.2_ region might prevent binding to the core enzyme since there would be no recognition of the β’ coiled-coil domain. However, mutagenesis of these residues in σ has shown that there is just a weakening in their binding to the coiled-coil domain [66]. In addition, thermal denaturation experiments indicate that mutant subunits are as stably folded as the wild types, suggesting that the principal effect of these mutations is the allosteric regulation of the regions σ_2.3_ and σ_2.4_, involved in melting and recognition of the −10 box [66]. On the other hand, all the endosymbionts have distinct variations in the σ_2.3_ region; however, all of them conserve the essential aromatic residues necessary for DNA melting and for the correct folding of the σ_2_ region (Fig 3a, violet sites).

The σ_2.4_ region presents exclusive substitutions of the residues involved in the recognition of the nucleotide in position −12 of the −10 box. Similar changes can be seen in all *Hodgkinia* strains, which conserve a histidine instead of glutamine in the position ECO: 437. Likewise, *C. Carsonella ruddii* PV, PC, HT, HC, DC, YCCR and BC replaced the ECO: T440 by a leucine or an isoleucine. Unlike the latter, *Tremblaya* strains and *T. phenacola* PAVE conserve these two sites (ECO Q437 and T440) (Fig 3a, orange sites). Previous studies using punctual mutagenesis in the region σ_2.4_ of *E. coli* σ^70^ (70 KDa) and *Bacillus subtilis* SigA (42.9 KDa), the latter is homologous to σ^70^ but smaller, indicated that changes in amino acids at these positions change affect specificity of each σ for its respective promoters [50]. In *E. coli* σ^70^ the substitutions in two independent residues ECO: Q437H and T440I were performed, and the results showed that σ conserves the capacity to recognize the nucleotide at −12, however, the specificity for promoters with a cytosine at this position was significantly higher [69]. Additionally, in SigA (the *B. subtilis* housekeeping σ equivalent to σ^70^ of *E. coli*) the replacement of ECO: Q196R (corresponding to ECO: Q437 in *E. coli)*, had a similar result as in *E. coli* [77].

Substitutions observed in the σ_2.4_ region might be related to variations in the specificity of σ for different promoters. According to previous studies in DNA-binding proteins, it has been observed that the substitution of amino acids such as lysine, asparagine, serine, methionine and aromatics like phenylalanine do not difficult their affinity to DNA [78]. However, these changes might exert mild effects on σ affecting its specificity for promoters. Based on structural predictions of σ factors for these endosymbionts, we observed structural similarities compared to *E. coli* σ^70^ (Fig 4). The changes observed in the σ_2.4_ region of *C. Hodgkinia* and *C. Nasuia* correspond to mutations already experimentally made in the *E. coli* σ^70^ and *Bacillus* SigA. Nevertheless, it is not possible to ensure the same specificity and recognition of the promoter on σ of endosymbionts because these are the result of a set of complex interactions between distinct regions of σ with other subunits of RNAP and with DNA promoters. So far, the results indicate that RNAP of *Hodgkinia* to exception of the Dsem strain, recognizes only the −10 box and the −10 extended promoter elements, because they lack the fragment required to form the β-flap domain that recognizes the −35 box (Fig 4 d, e, f and g).

Endosymbionts with reduced genomes also contain variable proportions of GC in their genomes. While *C. Carsonella* and *C. Nasuia* contain less than 18% GC, *C. Hodgkinia* and *C. Tremblaya* contain above 40%. In order to know if such variations of GC content in genomes correlate with changes in σ factors, we carried out a comparative analysis that included a phylogenetic tree. In this analysis we included (in addition to the 31 endosymbiotic bacteria) 13 homologues of σ^70^ coming from other endosymbionts and free-living bacteria with a lesser extent of genome reduction and containing similar, lesser or greater GC contents than the endosymbionts here studied (S2 Table).

In comparison to other endosymbionts with reduced genomes, we obtained that *T. phenacola* PAVE, princeps PCIT and PCVAL are the endosymbionts with the closest σ to *E. coli* σ^38^, instead of the canonical σ^70^ (S4 Fig). σ^38^ pertains to the σ^70^ family but in *E. coli* it is dedicated to transcribe stationary phase genes. Between these two kinds of σ (70 vs 38), there are some differences in the region σ_4.1_ that increase σ^38^ ability to bind to the β-flap domain region of the −35 box.

From this analysis, we were unable to determine any correlation between the GC content and changes in the σ factors. Regions of σ (such as σ^70^ from *E. coli*) are well preserved in all other bacteria, except for the case of *C. Zinderia insecticola CARI,* which is the organism with the lowest genomic GC content, but, unlike other endosymbionts with reduced genomes, it has a less preserved σ_4_ region and contains less substitutions in the σ_2_ region (Fig 3a).

### Studies on selective pressure of RNA polymerase subunits

To analyse the effect of amino acid substitutions observed in genes that encode for RNAP subunits, we made two types of selection analyses. Considering that bacterial endosymbionts are subject to an accelerated rate of molecular evolution [79], a useful tool towards determining the strength and type of selection acting on the genes that encode proteins is the estimation of the ratio of non-synonymous to synonymous substitutions (D_N_/D_S_) using phylogenetic codon-substitution models. Under neutral evolution, the rate of accumulation of D_N_/D_S_ should be 1. While a D_N_/D_S_ >1 is indicative of positive selection and a value <1 indicates negative selection [80]. We used this criterion to identify the kind of selection on genes that encode for the RNAP subunits from each group of endosymbiont bacteria (S3 Table).

#### Candidatus Hodgkinia cicadicola

The *Hodgkinia cicadicola* RNAP subunits reveal that *rpoB* and *rpoC* genes were subject to a purifying selection with lower D_N_/D_S_ values (0.3) in almost all strains, this might be because most of the nucleotide substitutions in these genes were synonymous. On the other hand, *rpoA* genes had a high purifying selection process in most *Hodgkinia* strains, while in Tetund 2, TETLON and TETMLI1 strains show neutral selection. Finally, the *rpoD* gene show an increased D_N_/D_S_ value, which means that the purifying selection is less rigorous in some *Hodgkinia* strains than in others, this lack of pressure results in greater diversity in protein regions, which possibly corresponds to the different coexistence conditions with other strains in each host (Fig 4).

#### Candidatus Tremblaya

For the strain PAVE, the *rpoA*, *rpoB, rpoC* and RpoD genes had ω values less than 0.02. Conversely, the strain PCVAL shows a neutral selection for all those same subunits (D_N_/D_S_>0.8). So, the relaxation of selective pressure is not the same for each strain of these bacteria (Fig 5).

**Fig 5.**
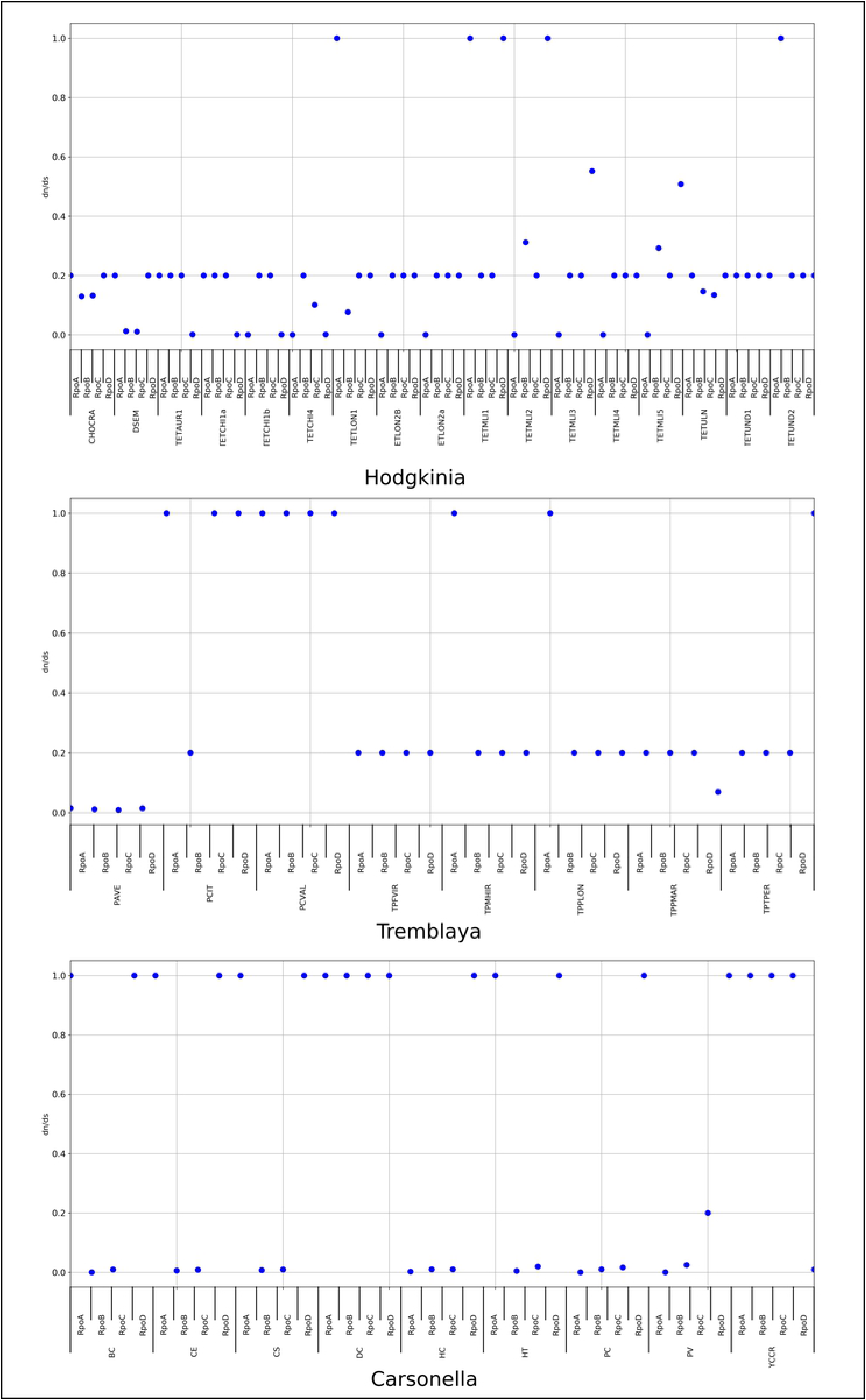
Selective pressure by branch analysis on the RNAP subunits in obligate endosymbionts. The relation D_N_/D_S_ for each of the subunits and for each endosymbiont was calculated. Here, we show the dispersion of the obtained values. Most genes are under negative selection, and in some cases, they display values greater than 1 (represented as 1.2), this does not mean that they are under positive selection (level of significance greater than 0.05), RNAP subunits shown a relaxed selection in all cases.

#### Candidatus Carsonella ruddii

Same as for the other endosymbionts, the *rpoB* and *rpoC* genes of *Carsonella ruddii* strains are more conserved than other subunits of RNAP, they had D_N_/D_S_ values less than 0.2 (except for DC and YCCR strains). Of these, the *rpoA* gene had a neutral selection in five strains (CE, CS, DC, HT and YCCR) and there is a strong purifying selection in the rest of *Carsonella* strains. On the contrary, the *rpoD* gene in all these strains had a neutral selection except for the BC strain, which might result in a σ factor containing variations in amino acid content (Fig 5).

Finally, we also performed a branch-site analysis for each subunit in each group of endosymbionts, where D_N_/D_S_ is constant among branches but varies among sites, this approach is used to detect punctual positive selection. Here, our aim was to identify nucleotides in the genes under positive selection. Positively selected amino acids were predicted by a Bayes Empirical Bayes (BEB) analysis for all the branches to find putative significant positive selection. The relationships among these positively selected amino acid residues were evaluated by mapping them on the respective three-dimensional protein structures to determine if these have functional implication (Fig 6).

**Fig 6.**
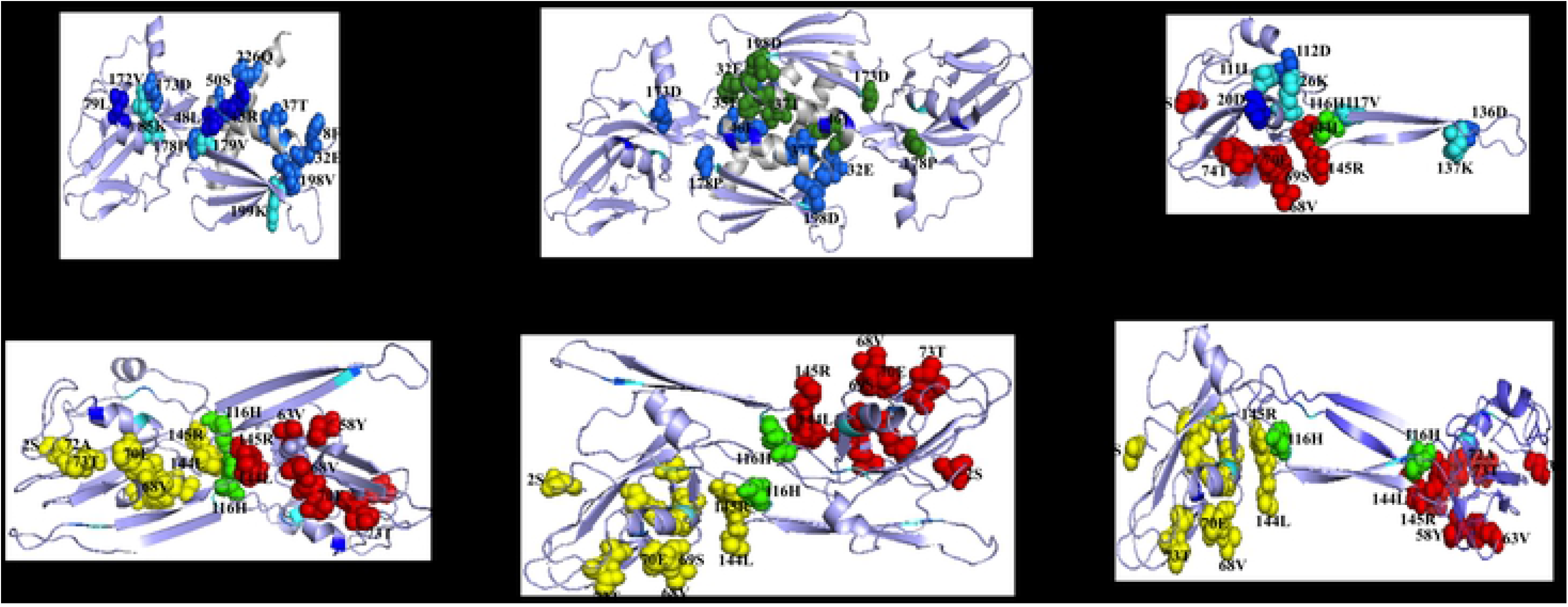
α-subunit structural comparison between *E. coli* and *C. Hodgkinia TETUND2*. 3D crystallographic structure of monomers and dimers of α-subunit in *E. coli* (4YPL) (a, b). 3D model of monomer α-subunit of *Hodgkinia* TETUND2, sites under positive selection are shown in red, H116 is shown in green (c). Predictions of α-subunit dimer formation in *C. Hodgkinia* TETUND2, sites under positive selection are shown in red and yellow in each monomer, H116 is shown in green (d, e, f). White regions in a and b structures are not conserved in the α-subunit of *C. Hodgkinia* TETUND2 (g). Amino acids in blue dark and turquoise are important for RNAP core formation. In the b structure, the sites in dark green are the same sites in blue in the structure a.

Positively selected amino acids were detected in the αNTD of *C. Hodgk*inia *Dsem* and TETUND2 (S4 Table). We mapped the selected amino acids with high level of support (BEB p>0.95) in the structural models of the α subunit of *Hodgkinia* TETUND2 and Dsem (Fig 6c, red amino acids and S5 Fig respectively). In the case of *C. Hodgkinia* Dsem the selected amino acids 55V, 95Q and 115H (S5 Fig) would be important for the correct folding of the αNTD. While, in *C. Hodgkinia* TETUND2 the amino acid residues 3A, 6K, 68E and 71T form a similar structure to that in the αNTD and, due to its localization, they could be involved in the interaction between the α subunit and both the β and β’ subunits (Fig 6c).

As previously mentioned, the *Hodgkinia* TETUND2 α subunit has lost part of the αNTD involved in the dimer formation. So, we evaluated if *Hodgkinia* TETUND2 α subunit can form the dimer, which is essential for RNA core formation [30]. For this, we used the software ClustalPro to model the α dimer, we obtained tree models with different size clusters of amino acids involved in dimer formation (Fig 6 d, e, and f). In these models, we mapped the sites under positive selection (Fig 6 d, e, and f, amino acids in red and yellow). Then, we performed *in silico* amino acid substitutions of the sites under positive selection, changing them by other amino acids present in the same sites of other *Hodgkinia* strains and *E. coli*. Finally, we used the software Bindx to evaluate the free-energy changes in protein-protein binding for the three models. We observed that the substitution of H116 by a proline in the three models changed the dimer formation in an energetically unfavourable way (S5 Table). Although the site H116 is not statically significant to be considered under positive selection, it would have an important role for stabilizing the dimer formation in this strain, however, experimental analysis is necessary to confirm this prediction.

## Conclusions

Endosymbionts are highly specialized organisms, which seems to be the result of co-evolution with their hosts. This might lead endosymbionts to evolve its genomic content to produce proteins that are simplified versions of their free-living counterparts. In this work, we studied the changes of RNAP subunits, trying to get an idea on how these enzymes carry out DNA transcription in these organisms.

We found that the β and β’ subunits tend to be conserved in all the studied endosymbionts, with just some differences in the regions involved in the interactions with the σ factor, possibly as result of major changes in σ. On the other side, the α subunit is more conserved in *C. ruddii* and *Nasuia* but than in other endosymbionts where the α subunit has lost its C-terminal domain. Furthermore, the total loss of the gene encoding this subunit has been reported in some strains of *Hodgkinia* [17], this may be due to complementation with genes from other bacteria, since these strains share host with other *Hodgkinia* strains. The strains Dsem, CHOCRA and TETULN are the unique strains known to not share their host with any other *Hodgkinia* strains and possibly that is why they preserve an α subunit, as we know for functional gene. On other hand, the rest of *Hodgkinia* strains share their host with one to five different strains, however, in these cases each *Hodgkinia* complex retains the α subunit genes [17]. We observed several amino acids under putative positive selection in the α subunit gene of *Hodgkinia* TETUND2, that would suggest a compensation in the loss of important regions for dimer formation in this subunit.

Some interesting events seem to be happening in all endosymbionts that do not occur in free-living bacteria. The σ factor encoding gene *rpoD* does not conserve the region 2, as does the conventional σ^70^ unit in *E. coli;* furthermore, we observed the loss of up to two domains (σ_1_ and σ_3_). Thus, drastic changes in endosymbiotic RNAPs coming from reduced genomes are related to sites and regions involved in promoter recognition, as we know in free-living organisms. These differences are not observed in any other free-living bacteria, not even in endosymbionts with larger genomes (*e.g*., with more than 170 Kpb genome size), which might suggest these changes as consequence of the drastic process of genome reduction that are experimenting these organisms.

In endosymbiont bacteria with reduced genomes it has not been possible to locate σ^70^ canonical promoters, not even for those the most conserved like those in ribosomal genes of bacteria [81], it might be, at least in part, due to the high A+T percentage in these genomes. However, *Hodgkinia* and *Tremblaya* have relatively high G+C% and lack intragenic regions, similarly to the rest of endosymbionts (more than 90% of their genome is composed of coding sequences) [6]. In addition, variations observed in σ factor regions are involved in the recognition of some promoter regions, hence, our results suggest that RNAP in endosymbionts conserve some capacity of sequence recognition, which is different to the σ^70^ consensus recognition sequence known for *E. coli*. However, experimental analysis must be done to corroborate this hypothesis.

Variations in recognition regions and sites of promoters might not be only ones in these bacteria. Previous reports indicate significant changes in the 16S ribosomal 3’ tail and its binding sequence, as well as in the Shine-Dalgarno element localized upstream of the protein-encoding genes [82].

Finally, we proposed a model of RNAP for each endosymbiont according to our findings (Fig 7). We observed that the RNAP of *Nasuia* conserved greater similarity to *E. coli* σ^70^ in comparison to other endosymbionts. According to this, it is expected that *Nasuia* RNAP recognizes the two promoters boxes (−10 and −35) and the two promoter accessory zones (−10 extended and UP element), in contrast to *Carsonella* RNAP, which seems to be unable to recognize the −10 extended box.

**Fig 7.**
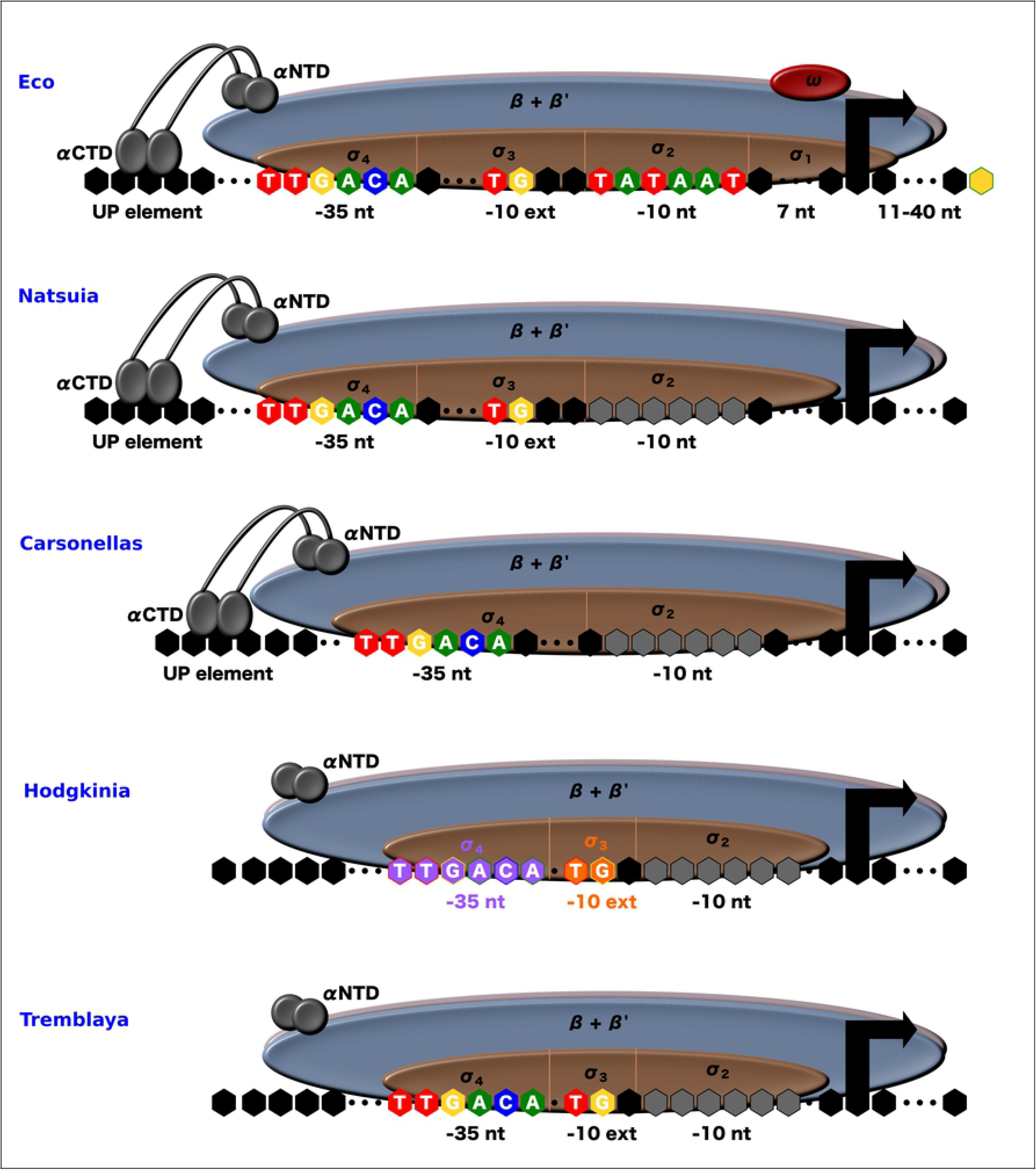
Proposed structural RNAP models of bacterial endosymbionts with reduced genomes. The conserved domains in each group of bacteria are identified in each model: *E. coli* RNAP model (Eco), *Natsuia* strains RNAP model, *Carsonella* strains RNAP model, *Hodgkinias* RNAP model (In orange, the TETULN, TETUND1 and TETUND1 strains exclusively using −10 extended region. In purple, Dsem strains exclusively using the −35 element), and *T. phenacola* PAVE, princeps PCIT and PCVAL RNAP model (Fig based on Busby and 2004 and Rangel-Chavez et al, 2019).

On the other hand, in the case of *Tremblaya* and *Hodgkinia* RNAP, due the absence of α C-terminal domain, they are unable of recognizing the UP element, and in the case of *C. Hodgkinia Dsem* neither the −10 extended promoter element. Contrary to *C. Hodgkinia Dsem*, other *Hodgkinia* strains are expected to recognize the −10 extended promoter element and, in this way, might compensate the loss of the β-flap-helix domain. The model of RNAP proposed for *Hodgkinia* suggests that it no longer conserves the mechanisms for transcription initiation common in free-living bacteria, as they lack the β flap-helix domain. In free-living bacteria, the β flap-helix domain is essential for the assembly of sigma to the RNAP core.

In a whole view, and regarding the study of RNAP subunits, we conclude that these four groups of endosymbionts with extreme-genome reduction do not follow the same evolutionary path, and that each strain has particular modifications, which may be a consequence of the conditions exerted by the host and/or the coexistence with other endosymbionts.

## Acknowledgments

CPR-C has a PhD fellowship (380338) from CONACYT. México. Authors thank Rafael Montiel for guiding on the selective pressure analyses and to Diego Andrés López Castro and Paola Angulo-Bejarano for reading the manuscript and to anonymous referees for useful comments to the document.

## Author Contributions

Conceptualization: AM-A and CPR-C

Formal analysis: Cynthia Paola Rangel-Chávez, Edgardo Galán-Vásquez, Azucena Pescador-Tapia.

Funding acquisition: Agustino Martínez-Antonio.

Supervision: Agustino Martínez-Antonio.

Writing – original draft: Cynthia Paola Rangel-Chávez, Edgardo Galán-Vásquez, Agustino Martínez-Antonio.

Writing – review & editing: Agustino Martínez-Antonio, Luis Delaye Arredondo.

## Supplementary material

S1 Table. Complete list of studied endosymbionts

S2 Table. Bacteria with similar %GC to endosymbiotic bacteria with reduced genomes

S3 Table. Results obtained of selective pressure analysis by branch model

S4 Table. Results of selective pressure obtained by branch-site model

S5 Table. Changes of free energy in selected sites of α subunit homodimer

S1 Fig. Interaction domains in α subunit alignment

S2 Fig. Interaction domains in the β subunit alignment

S3 Fig. Functional domains and sites in β’ subunit alignment

S4 Fig. Phylogenetic Tree of sigma 70 between endosymbionts with extreme genome reduction and other bacteria

S5 Fig. Selected sites mapped in *Hodgkinia* Dsem α subunit

